# Two particle picking procedures for filamentous proteins: SPHIRE-crYOLO filament mode and SPHIRE-STRIPER

**DOI:** 10.1101/2020.02.28.969196

**Authors:** Thorsten Wagner, Luca Lusnig, Sabrina Pospich, Markus Stabrin, Fabian Schönfeld, Stefan Raunser

## Abstract

Structure determination of filamentous molecular complexes involves the selection of filaments from cryo-EM micrographs. The automatic selection of helical specimens is particularly difficult and thus many challenging samples with issues such as contamination or aggregation are still manually picked. Here we present two approaches for selecting filamentous complexes: one uses a trained deep neural network to identify the filaments and is integrated in SPHIRE-crYOLO, the other one, called SPHIRE-STRIPER, is based on a classical line detection approach. The advantage of the crYOLO based procedure is that it accurately performs on very challenging data sets and selects filaments with high accuracy. Although STRIPER is less precise, the user benefits from less intervention, since in contrast to crYOLO, STRIPER does not require training. We evaluate the performance of both procedures on tobacco mosaic virus and filamentous F-actin data sets to demonstrate the robustness of each method.

## 1. Introduction

The determination of protein structures using single-particle cryo-EM requires the selection of thousands of particles within micrographs. Various methods have been developed to automate this task [1–9]. Especially the introduction of deep learning based procedures have dramatically reduced the false-positive rates of picking and made automatic picking of particles the standard in single particle cryo-EM [5–9]. The picking of filaments is more challenging because of the line-like structure of the specimens. It is especially difficult to omit filament crossings and overlaps. Although procedures have been introduced that allow the automated picking of helical samples [10,11], a deep learning based helical specimen picker is missing.

Here, we present a new deep learning filament picking procedure implemented in our single particle selection tool SPHIRE-crYOLO [5]. CrYOLO is based on a convolutional neural network (CNN) and the You Only Look Once (YOLO) approach [12]. CNN is a deep learning network architecture that became prominent in machine learning during the last ten years. Today, CNNs are the state-of-the-art choice for image classification and object localization.

A traditional CNN-based classifier trained on a set of positive (e.g. particles) and negative examples (e.g. contamination or background) can be turned into an object detection system by using a sliding window. This moving window, slides over the input image, crops out small regions from it and then classifies those regions as either containing a particle or not. This allows the localization of particles within micrographs. However, this approach has very limited spatial contextual information and is slowed down by a high computational overhead.

The You Only Look Once (YOLO) framework by Redmon et al. [13] is an alternative to the sliding window approach: Instead of many small cropped out regions, the whole micrograph goes through the network in a single pass. Internally, the image is divided into a grid where each grid cell is responsible for predicting a single box. The confidence that a grid cell actually contains a particle, the relative box position inside the grid cell and the width and height of a box is estimated by each individual grid cell. This approach reduces the computational overhead and makes YOLO fast while retaining its accuracy. Moreover, because the network sees the complete image at once it is also able to learn about the spatial context of particles. These properties make the generic YOLO framework an excellent basis for particle picking in crYOLO. CrYOLO enables the automated picking of particles within low signal-to-noise ratio cryo-EM micrographs with minimal human supervision or intervention.

In the new filament mode, crYOLO places boxes on the filaments after training on several manually labeled micrographs. An extra post-processing step uses these boxes as support points to trace the actual filaments. As crYOLO always takes the larger context into account, it is able to skip dense filament regions, or broken areas sections of the filaments without the need for additional, user selected thresholds. This enables crYOLO to identify filaments on previously unseen micrographs, with an accuracy that is similar to manual picking.

In addition, we present STRIPER as an alternative to the filament mode in crYOLO. STRIPER enhances linear structures within in an image using oriented Gaussian smoothing kernels and then applies a line detecting-algorithm. Potential crossings are detected by the same algorithm and can be skipped.

## 2. Material and Methods

### 2.1 Oriented Gaussian filtering for feature extraction

An oriented Gaussian smoothing kernel can be used to extract direction dependent information from an image and/or enhance specific directional features of an image. Here we use an oriented Gaussian smoothing kernel to extract line features. The second derivative in the y-direction *M*(*x, y*) of a smoothing kernel is given by:

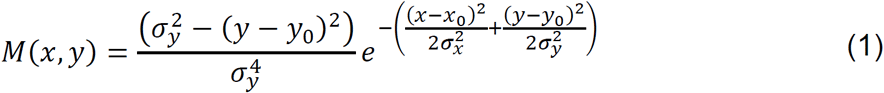

Where *σ*_*x*_ and *σ*_*y*_ are the spread in the x- and y-direction, respectively, and *x*_0_and *y*_0_ denote the center of the mask. The spread *σ*_*y*_ determines the amount of averaging in the y-direction and the spread *σ*_*x*_ is proportional to the width of the line-structure it enhances (see mask in Fig. 1).

**Figure 1.**
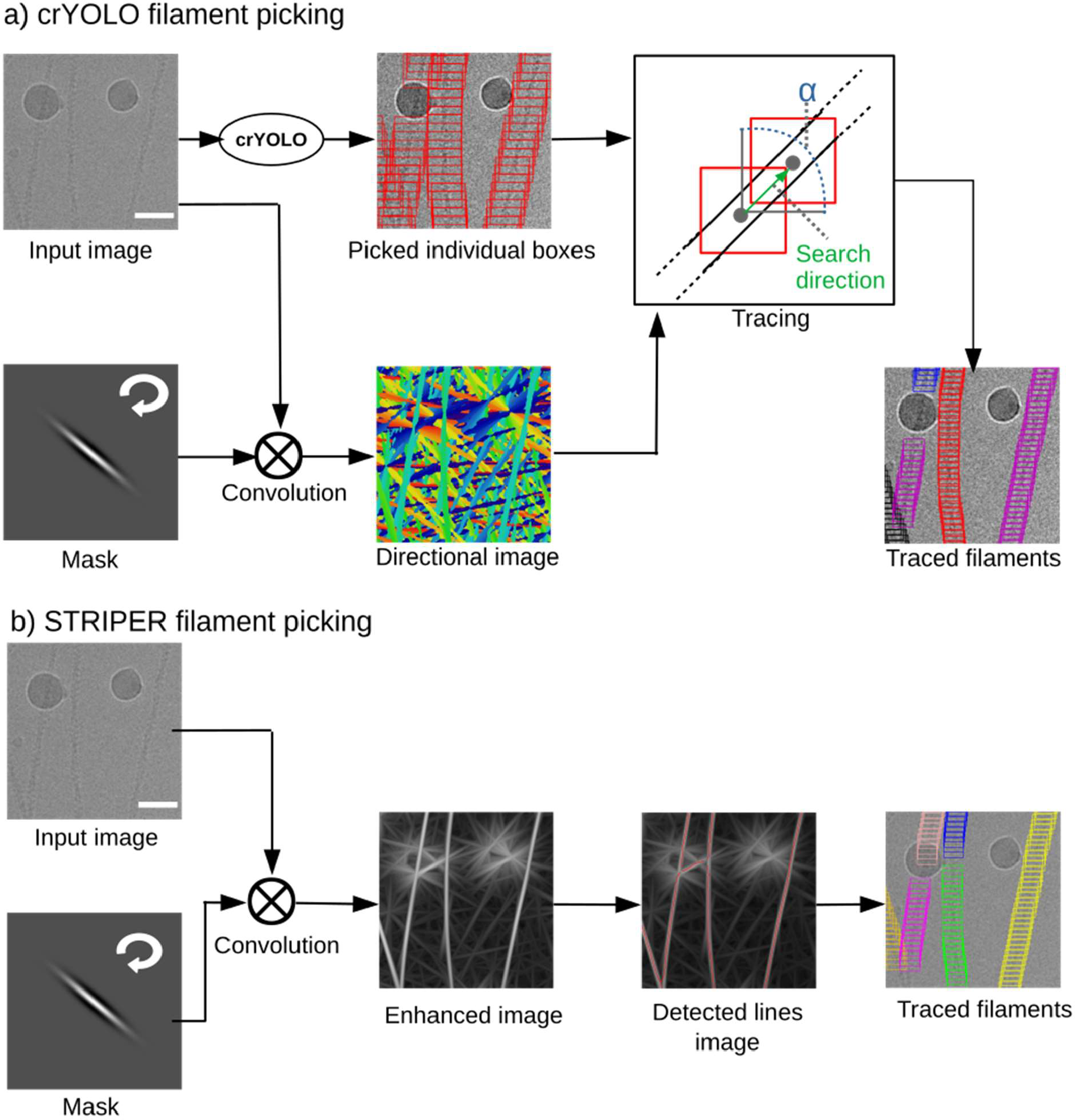
Filament picking with crYOLO and STRIPER. (a) CrYOLO: The input image is convolved multiple times with rotated versions of the convolutional mask. Each pixel in the directional image is color-coded to indicate the direction of the mask with the strongest response at its coordinates. During tracing, a box is randomly chosen and the search direction is determined by the directional image. The search angle alpha is set to 120°. The search radius is set proportional to the box size. Finally, given traced boxes the filament boxes are generated using a pre-set distance. (b) STRIPER convolves the input image with the same mask as crYOLO. An enhanced image is created by setting each pixel value to the strongest response of the rotated convolutional filters. The enhanced lines are then detected by a line tracing algorithm *[14,20]*. After tracing, crossing points are removed and the boxes are placed along the detected lines in a user defined distance. Scale bars, 50 nm.

To enhance lines with a specific orientation *θ*, M needs to be rotated by *θ*. Let *I(x,z)* be our input image; then the oriented filter response *R(x,y)* enhances structures that are oriented in the direction *θ*:

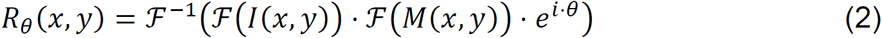

where *ℱ* denotes the Fourier transform. To enhance all line structures with arbitrary orientation, we calculate:

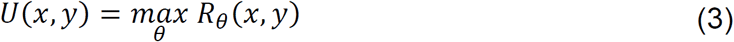

To extract the dominating direction at every position in an image, we calculate:

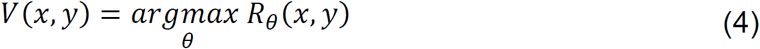

*V* is used in the crYOLO tracing method, and *U* is used in STRIPER for running the line detection algorithm.

### 2.2 Steger line detection

The STRIPER ridge detection algorithm is based on [14]. It identifies any lines present within the image through differential geometric properties. More precisely, the algorithm is divided into four steps:

1. Pre-processing based on Koller’s approach [15] where the image is filtered with the derivative of a Gaussian smoothing kernel. The resulting image features a series of mathematical properties [15] that allow the algorithm to detect lines of arbitrary width.
2. Detect all the pixels on an identified line segment (‘line points’). For each line point a strength *s* is calculated which is a measure of belonging to a particular line. In a greyscale image, pixels that are not line points are assigned an *s* value of 0.0, while line points closer to the center of a line have an *s* value of up to 1.0.
3. Connect line points to form the actual lines and identify line crossings. The list of line points *L* is first reduced by removing any line points with an *s* value lower than a user defined threshold. The procedure for building a generic line *o* is a generalization of a hysteresis threshold operation [16]:
  a. Select line point *p* with the highest *s* value as the starting point of new line *o*.
  b. From the surrounding pixels of p, select the one with highest *s* value that is not already part of *o* and add it to *o*.
  c. Repeat (b) until:
    i. No valid line point is found, thus indicating the end of the line.
    ii. The selected line point is already part of a different line. Mark this point as a junction and split the line in two. New lines are created until all points in *L* have been visited once [14].
4. Determine line width. Since the edges of a thick line are lines themselves, they are identified in a similar way as above by using a different filter.

Steger uses a generic line model defined by the *s* values of each pixel in step **(2)**, and the computed line widths, to improve the position of the estimated line [14].

### 2.3 crYOLO

CrYOLO is a particle picking procedure based on the YOLO framework [12]. For a technical description of crYOLO we refer the reader to our original publication [5].

### 2.4 Evaluation procedures

For the evaluation of the proposed procedures we used the common metrics of recall and precision. The recall score measures the ability of the classifier to detect positive examples, and the precision score measures the ability of the classifier to not label a true negative as a true positive. Both measurements are commonly used for binary classification tasks. To calculate the precision recall for the user selected filaments *T*_*u*_ and the results *T*_*p*_ given by crYOLO or STRIPER we did the following:

1. We transformed traced filaments of *T*_*u*_ into a binary image *B*_*u*_ by setting all pixels along the filament and within a local a radius of *f*_*w*_/3 to the value 1. The same is done for *T*_*p*_ which results in the binary image *B*_*p*_.
2. We calculated a difference image *D* by *D* = *B*_*u*_ − *B*_*p*_. Here we define *D*_1_ as binary image where all pixels in D are set to 0 except the positive pixels and *D*_−1_ where all pixels in D are set to 0 except the pixels with negative values.
3. The false-negative pixels (FN) and the false-positive pixels (FP) are then given by

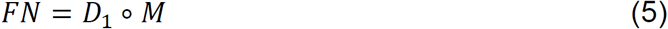

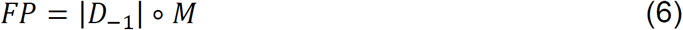

Where M is a mask of ones of size *f*_*w*_ and ∘ the morphological opening operator. The true positive pixels are then given by

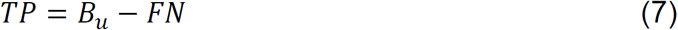

Finally, the precision and recall are defined as

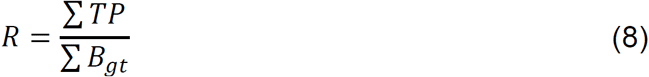

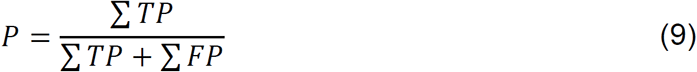

To calculate these statistics, we ignored both picks on the border of the image and the start and end positions of each filament as they are connected with high uncertainty during manual selection.

## 3. Results and discussion

### 3.1 CrYOLO filament mode

Since we have just recently introduced the crYOLO filament mode, a generalized model is not available yet. Therefore, crYOLO requires several filaments to be labeled manually in order to properly train the model. We used the e2helixboxer program provided by EMAN2 [17] for manual selection, but any other program that allows manual picking of filaments can be applied. The number of micrographs that need to be manually annotated might vary dependent on the filament density, orientation and background variations. Also a project with many aggregates might increase the number of training images needed to get a good working model.

The training of crYOLO works as described for single particle projects [5] : The network is trained on the manually labeled micrographs, while a small subset of those micrographs is used for validation. After each round of training, crYOLO measures the success of picking on the validation micrographs and stops the training when the validation performance does not increase anymore. After training is completed, crYOLO goes through the data set and places boxes on filaments. The boxes have no defined distance to each other and no information about to which filament they belong. As a special requirement for helical specimens, the filament mode in crYOLO allows more overlap of boxes during picking. The originally determined positions of the boxes on the filaments are merely used as support points for placing new boxes with a user defined distance. During this postprocessing step, an oriented convolutional mask filter is used to estimate the direction of every filament.

The mask contains the second derivative of an oriented Gaussian smoothing kernel (see Eq. 1). Given the user-selected filament width *f*_*w*_ in pixels and a default mask width *m*_*w*_ of 100 pixels, then *σ*_*x*_ and *σ*_*y*_ are defined using the “full width at half maximum” criterion:

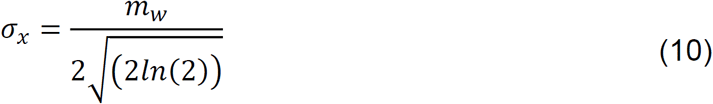

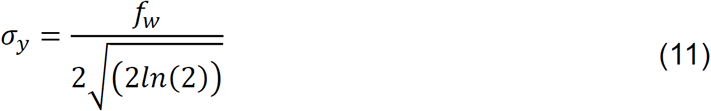

The kernel is rotated and each rotated version is convolved with the input image. For each pixel in the input image, the rotational angle of the convolutional mask that gives the maximum response is determined (see Eq. 4), which, evaluated for all pixels, gives the directional map (Fig. 1a).

Next, a box is chosen randomly on the filament and the direction of the filament is evaluated by the directional map. The next box is searched for within a radius proportional to the box size; the search is restricted by the estimated direction of the filament. If a box can be found, the search is continued. If not, another yet untraced box is selected, and the search is repeated. In case the search finds a previously traced box, both filament segments will be merged into a single filament if their directions are comparable. After the tracing is done, new boxes are generated based on the user defined distance. The final boxes are saved in STAR and EMAN helical box format.

The filament mode of crYOLO has already been successfully applied for solving the structure of F-actin in complex with drug-like toxins [18] and Lifeact [19].

### 3.2 STRIPER

Instead of using a trained deep neural network to identify the boxes along the filaments, the STRIPER filament picking procedure is based on a classical line detection approach (Fig. 1b). Cryo-EM images typically have a very low signal-to-noise ratio which is problematic for line detection algorithms. We therefore included a line-enhancing pre-processing step in STRIPER. In this step, the filaments are enhanced using the same oriented Gaussian smoothing kernel as described above for crYOLO. The width of the mask is configured by the user and should be set to the filament width. The lines in the enhanced image (see Eq. 3) are then extracted by the ridge detection algorithm of Steger [14,20].

To run the STRIPER filament procedure, four parameters need to be provided: (a) The filament width in pixel, which can easily be measured; (b) the mask width in pixel, which is set to 100 by default, and only has to be changed for very flexible filaments (both parameters are used for the creation of the enhanced line image); (c) the upper and (d) lower threshold used as hysteresis thresholds for the Steger’s line detection algorithm (see Materials and Methods). Since the latter values are difficult to guess, STRIPER provides an optimization algorithm to estimate these parameters. For this, the user has to manually select the filaments in two to three micrographs. While fixing the lower threshold to a value of 0, a simple grid search will then find the best upper threshold to detect as many of the annotated filaments as possible. Finally, the lower threshold is increased stepwise to remove false-positive detections. After extracting the lines, STRIPER splits them at crossing points and boxes are placed along the lines with a user defined distance.

By defining a minimum length of detected filaments, STRIPER can remove short line-like contaminations that are detected as false-positives. Moreover, STRIPER allows the user to provide a binary mask for each micrograph. This mask divides an image into valid picking regions and regions with carbon or contamination. This masking option is especially useful as deep learning-based carbon and contamination detection just recently became available [7,21]. These programs determine valid and non-valid regions with high accuracy and create binary masks, which then can be directly used in STRIPER to remove false-positive selections.

STRIPER has already been successfully applied for solving the structure of toxin-stabilized F-actin [18] and F-actin in the ADP-P_*i*_ state [22].

### 3.3 Training and configuration

To test both procedures we used the publicly available tobacco mosaic virus (TMV) data set (EMPIAR 10022) [23] and one of our in-house F-actin data sets [19]. The TMV data set has two difficulties: First, several filaments are localized on the grid in very close proximity and should ideally not be selected. Second, the TMVs contain interruptions or discontinuities in their structure, which should be excluded from selection. The challenge of the F-actin data set is that often filaments are crossing each other and that large carbon areas are covered with F-actin. In contrast to the TMV data set, the F-actin images contain carbon edges, which is especially demanding for the selection process, since they appear as line-like structures.

To train crYOLO on both data sets, we used manually labeled filaments on several micrographs. For F-actin, we selected 275 filaments on 24 micrographs. For TMV, we used the manually traced filaments that were provided with the EMPIAR data set and selected a subset of 425 labeled filaments from 44 micrographs. We roughly estimated the width of TMV and actin filaments on the images and used these values for processing (∼200 Å for TMV and ∼ 80 Å for F-actin). Since the distance of the boxes is not relevant for the selection of the filaments, we used a standard box distance of 20 pixels for both data sets. When processing the data further for structural investigations, this value should be adjusted, taking the helical rise of the filament into account. The picking threshold in crYOLO was set to the default value of 0.3 for both data sets. For evaluation, 20% of the labeled micrographs were not used during training.

In STRIPER, several processes that are automatically done in crYOLO, such as binning, normalization and filtering have to be done manually. Thus for testing STRIPER, we binned the TMV and F-actin images by a factor of 4, low-pass filtered them with an absolute cut-off frequency of 0.1, and normalized them by subtracting the mean and dividing by the standard deviation. All pixel values greater than 3 or lower than -3 were saturated.

The contrast of an image has a strong influence on the line detection algorithm used in STRIPER. Since the contrast depends very much on the defocus at which the images have been taken, we manually labeled filaments in one micrograph with high defocus and one micrograph with low defocus to determine the hysteresis thresholds. Then, we applied the internal grid optimization routine to find the best set of selection thresholds (upper and lower threshold of 0.77 and 0.29 for F-actin and 0.2 and 0.1 for TMV, respectively).

For the evaluation of precision and recall for crYOLO filament mode and STRIPER we used the same micrographs.

### 3.4 Evaluation on test data sets

When we applied crYOLO to the TMV data set, the recall and precision was 0.98 and 0.84, respectively. The automatic picking of filaments resulted in a selection of filaments that was comparable to the manually picked data sets. Filaments that touched each other were skipped and discontinuities in the filaments, which are typical for this TMV data set (Supplementary Fig. S1), were mostly omitted (Fig. 2g). In contrast, STRIPER identified and picked almost all filaments on the micrographs (Fig. 2e). This led to a high recall (1.0) but, since STRIPER also selected filaments that sit very close to each other and contain discontinuities, the precision of only 0.52 was quite low.

**Figure 2.**
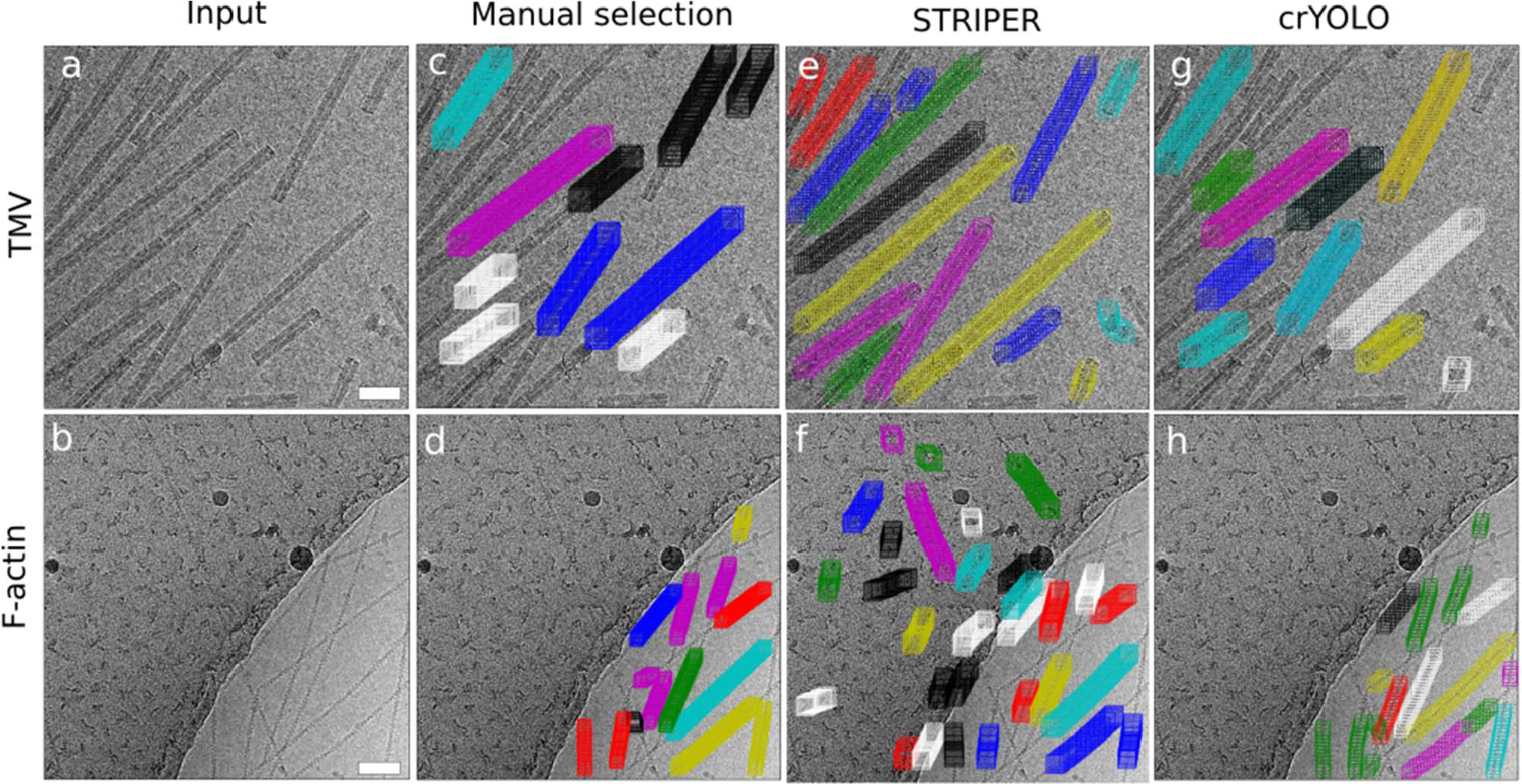
CrYOLO and STRIPER evaluated on micrographs with F-actin and TMV. (a,b) Input images that were not used during training (crYOLO) or parameter optimization (STRIPER). (c,d) Manually selected filaments. (e-h) Automatically selected filaments by STRIPER (e,f) or crYOLO (g,h). Scale bars, 50 nm.

For F-actin filaments, crYOLO achieved a recall of 0.95 and a precision of 0.83. It skipped most of the filament crossings and did not select the carbon edge or filaments on the carbon (Fig. 2h). STRIPER also identified most of the filaments and skipped their crossings (Fig. 2f). However, it also picked the carbon edge as well as filaments and line-like contaminations on the carbon. The recall was 0.81 and precision 0.51. As STRIPER supports binary masks, we masked out the carbon area using the MicrographCleaner tool [21] and repeated the selection (Supplementary Fig. S2). The masking led to an increase in the precision of STRIPER to 0.73 while the recall remained at 0.81.

We further assessed the quality of the selected filaments by 2D classification in SPHIRE [24] [25]. We then applied Cinderella [26], a deep learning based tool trained to identify high quality particle classes and used the percentage of rejected classes as indication for the quality of the picking procedure.

The particles picked by both selection procedures resulted in many high quality class averages (Fig. 3). Almost all classes calculated for the filaments identified by crYOLO were accepted by Cinderella (Table 1, Supplementary Fig. S3 and S4), demonstrating that crY-OLO indeed did not pick background, contamination or carbon edges.

**Table 1:** Results of 2D classification for F-actin and TMV. The number of class averages, and the number of classes accepted by Cinderella were evaluated for F-actin and TMV for crYOLO filament mode, STRIPER and STRIPER with masks.

| Dataset | Procedure | Accounted particles | Unaccounted particles | Total classes | Accepted classes |
| --- | --- | --- | --- | --- | --- |
| F-actin | crYOLO | 102,262 | 175 | 1,023 | 1,019 |
| F-actin | STRIPER | 112,118 | 897 | 1,128 | 708 |
| F-actin | STRIPER (+masks) | 69,671 | 217 | 697 | 682 |
| TMV | crYOLO | 13,338 | 154 | 133 | 133 |
| TMV | STRIPER | 49,897 | 1,333 | 503 | 496 |

**Figure 3.**
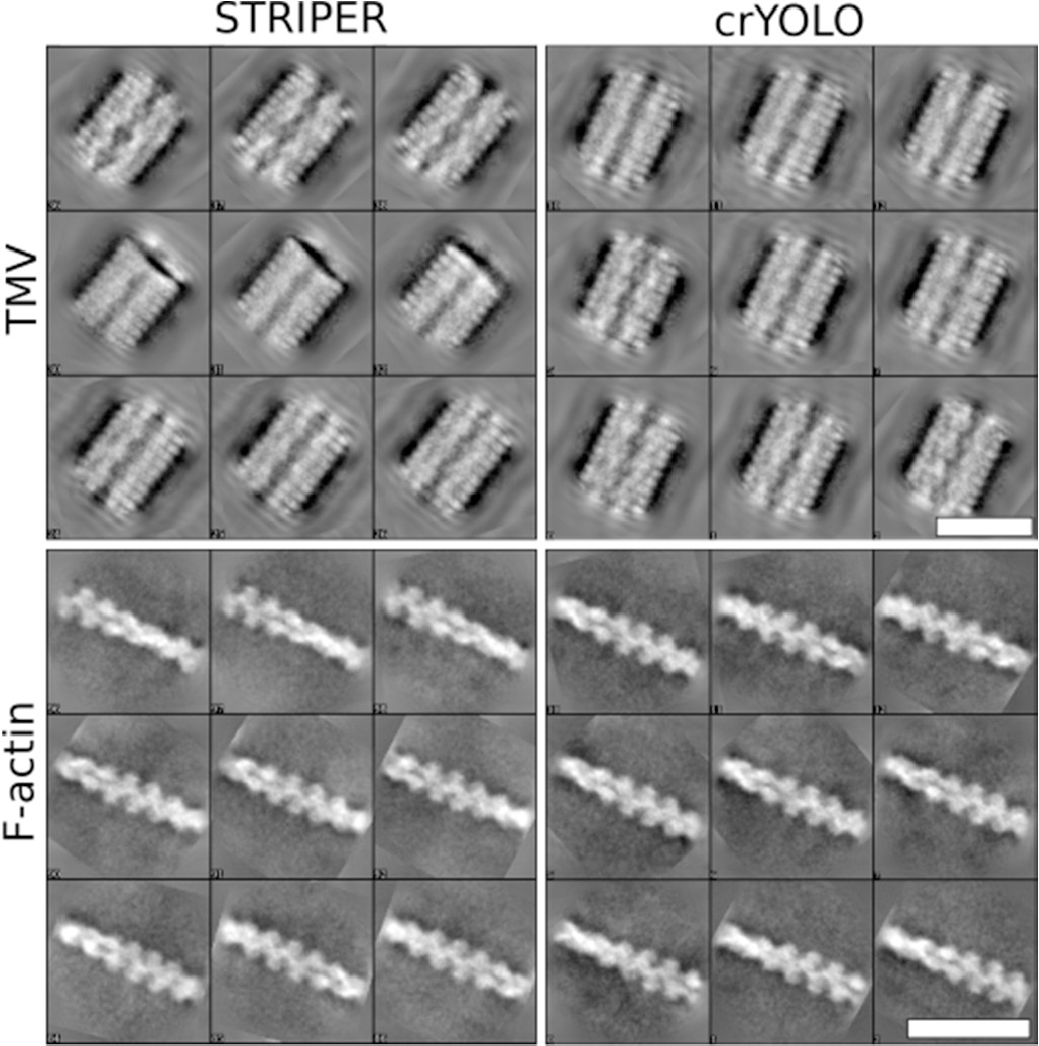
Example class averages calculated in SPHIRE. F-actin and TMV were picked by crYOLO filament mode or STRIPER. The respective number of class averages is listed in Table 1. Scale bars, 25 nm.

For particle stacks selected with STRIPER, almost all class averages were accepted by Cinderella in the case of TMV (Supplementary Fig. S5). The number of selected particles is much higher than for crYOLO, because STRIPER picked also TMVs with discontinuities and filaments that were in close proximity to their neighbors. As expected, 37% of the classes mainly contained false-positives and were rejected in the case of F-actin (Supplementary Fig. S6). Applying a mask to exclude the carbon and carbon edges solved this problem and almost all classes were accepted by Cinderella (Table 1). However, the total number of accepted classes is much lower compared to the classes obtained from particles selected by the crYOLO filament mode (Table 1).

### 3.5 Computational efficiency

To determine speed we picked 203 F-actin micrographs (4096×4096) using an Intel Xeon Gold 5122 CPU and a Nvidia Titan V GPU (32GB RAM). On average, crYOLO picked a micrograph in 7 seconds (including filtering and filament post-processing), while STRIPER required 0.8 seconds (without filtering). In both cases the algorithms presented in this paper provide highly efficient picking methods that allow users to pick large data sets in short amounts of time.

## 4. Conclusions

Particle picking is a crucial step in the cryo-EM processing pipeline. Picking of helical specimens is particularly challenging as crossings, overlaps and filaments in too close proximity to each other need to be omitted. This challenge is well illustrated by the data sets used as examples in this work. Here we present two picking procedures that allow accurate picking of filaments even for complex data sets as illustrated for TMV and F-actin in this work. One of them is based on the deep learning particle picking procedure crYOLO and the other, called STRIPER, is based on a line detection algorithm. CrYOLO is to our knowledge the first deep learning based particle picking procedure which supports filaments. Through this approach crYOLO learns the spatial context of filaments enabling it to omit carbon edges and too dense or broken filaments, thereby accurately reproducing the accuracy of manual picking. As there is no general model for filaments available yet, crYOLO requires training on manually selected micrographs. In contrast, STRIPER only requires four parameters which can be quickly determined by the integrated optimization procedure. While this makes STRIPER a very fast, easily accessible picker, it comes at the cost of reduced precision compared to crYOLO. However, we also showed that the usage of a binary mask significantly improves the precision of STRIPER, resulting in high-quality 2D classes. Both picking procedures produce box files that are compatible with the majority of cryo-EM processing software and come with standard hardware requirements. Considering the performance and accessibility, we believe that both procedures are a seminal contribution to the cryo-EM field.

## Acknowledgements

The authors thank T. Shaik for carefully reading the manuscript and valuable comments.

## Data availability

All data supporting the findings of this study are available from the corresponding author on reasonable request.

## Author contribution

Conceptualization; T.W. and S.R.; Software - crYOLO: T.W.; Software – STRIPER: T.W., L.L.; Software – Testing: T.W., S.P., F.S. and M.S.; Formal Analysis: T.W.; Writing – Original Draft: T.W. and S.R.; Writing – Review & Editing: S.P., T.W., S.R., F.S.; Funding Acquisition: S.R.

## Code availability

The filament mode of crYOLO has been available since crYOLO version 1.3. CrYOLO itself is free for academic use and can be downloaded together with the source code at http://sphire.mpg.de/. The version of STRIPER used in this paper was first implemented as an ImageJ [27] plugin and is open source. The source code can be found https://github.com/MPI-Dortmund/ij_striper. A Python implementation of STRIPER is currently being implemented and the alpha version can be found https://github.com/MPI-Dortmund/striper.

## Supplementary figures

**Supplementary Figure 1.**
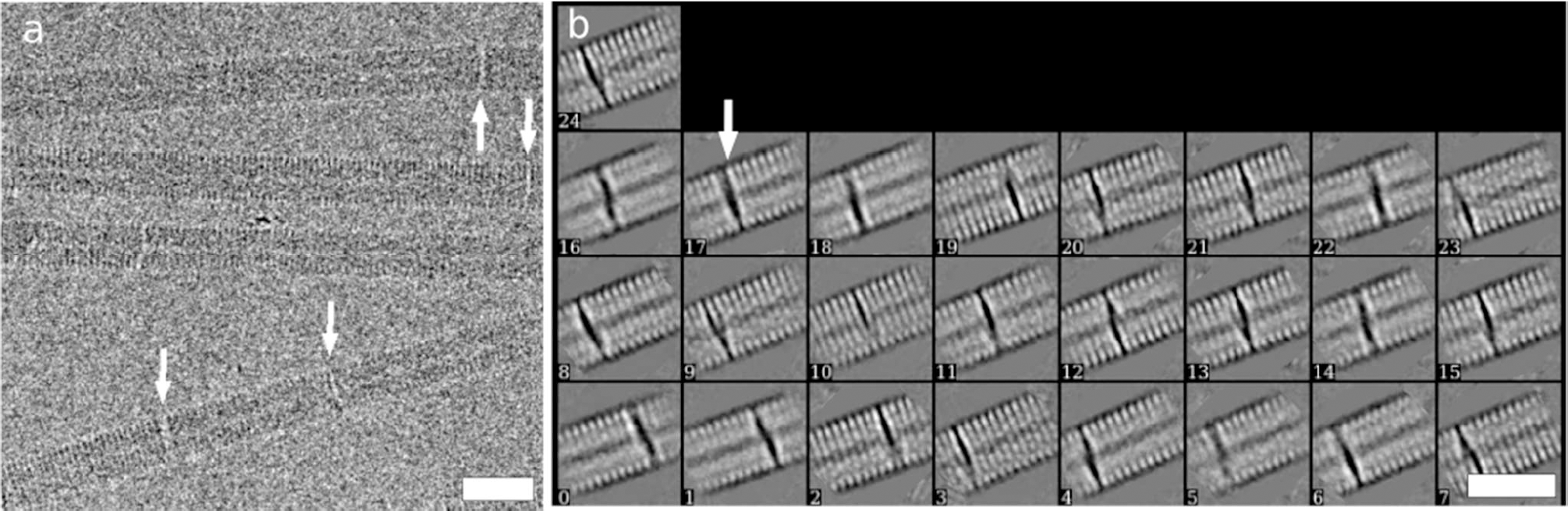
Discontinuities in TMV. (a,b) After picking with STRIPER (a) and 2D classification with SPHIRE discontinuities in TMV (highlighted by white arrows) also appear in 25 out of 503 classes (b). For crYOLO, only 1 class out of 133 shows a discontinuity. Scale bars, 25 nm.

**Supplementary Figure 2.**
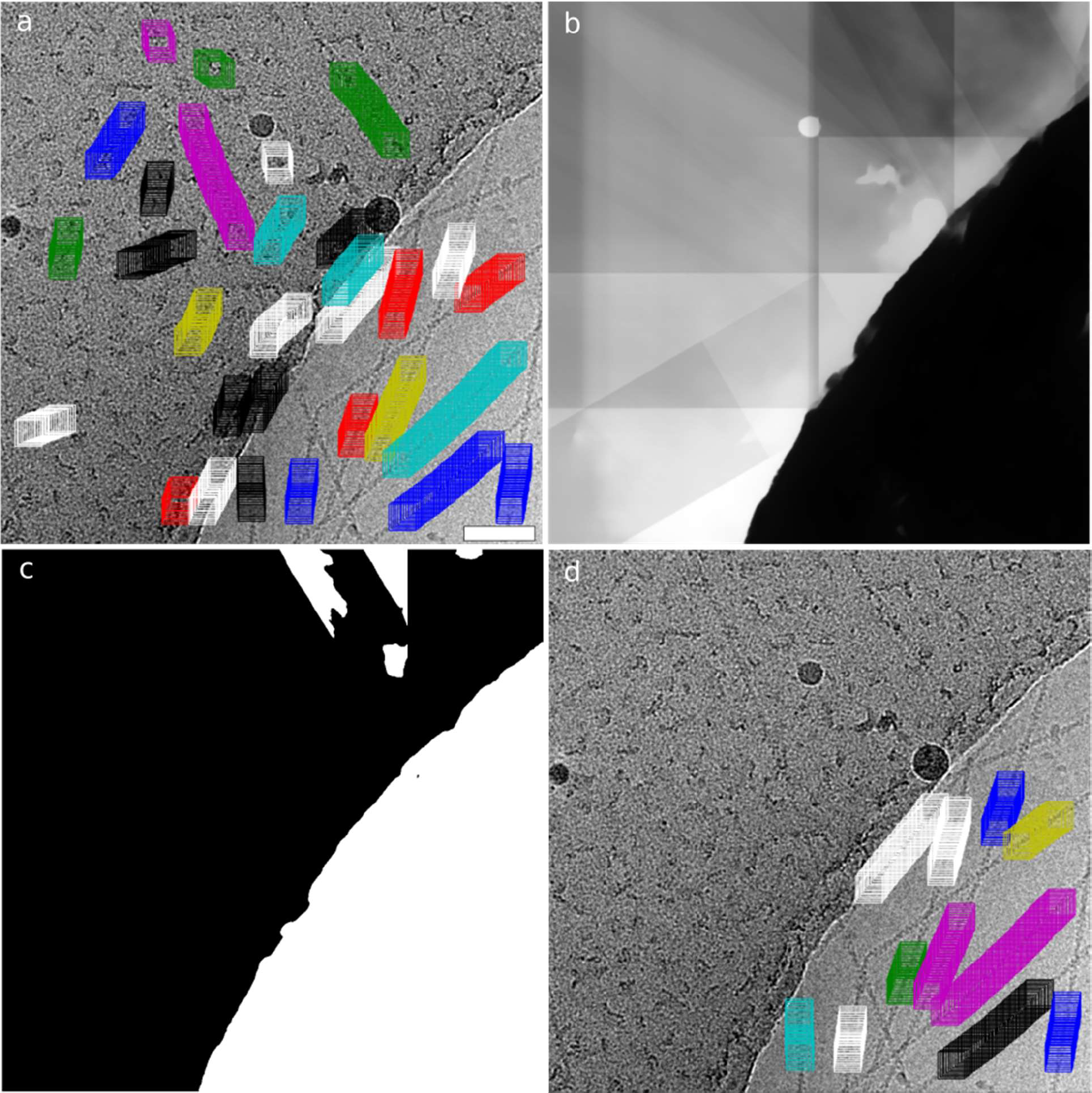
Elimination of false-positive STRIPER picks using a mask. (a) STRIPER tends to identify all line-like structures. This is problematic in case of line-like contamination or filaments on carbon. (b,c) Using MicrographCleaner we calculated a mask (b) and binarized it using a threshold of 0.3 (c). (d) This mask can then be used in STRIPER to remove false-positive filament picks. Scale bar, 50 nm.

**Supplementary Figure 3.**
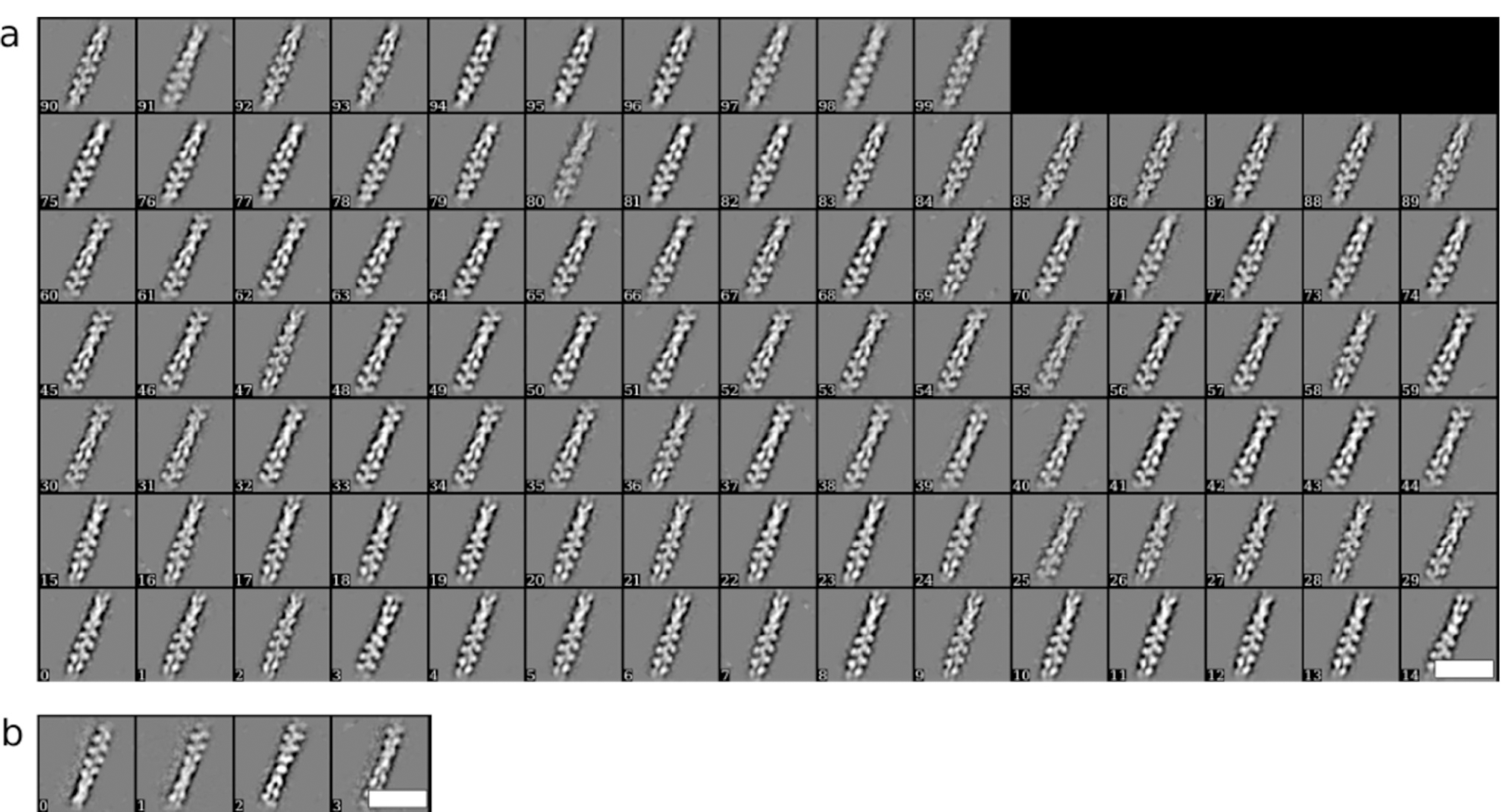
2D classes of F-actin accepted or rejected by Cinderella. (a) 100 out of 1023 accepted classes and (b) all rejected classes for filaments picked by crYOLO. Note that the rejected classes seem to be false-positives. Scale bars, 25 nm.

**Supplementary Figure 4.**
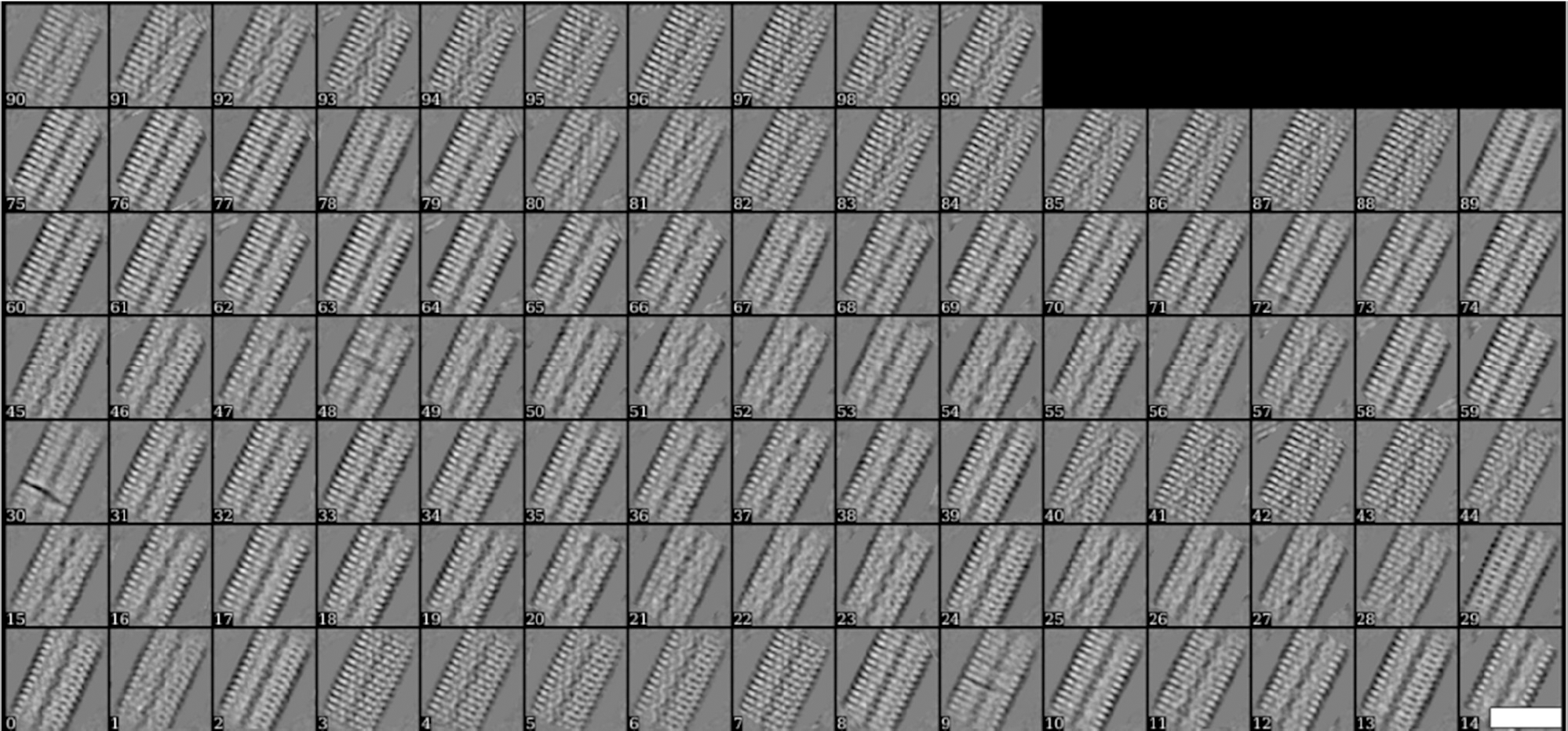
2D classes of TMV filaments accepted by Cinderella. 100 out of 113 classes accepted by Cinderella. Filaments were picked by crYOLO. No class was rejected by Cinderella. Scale bar, 25 nm.

**Supplementary Figure 5.**
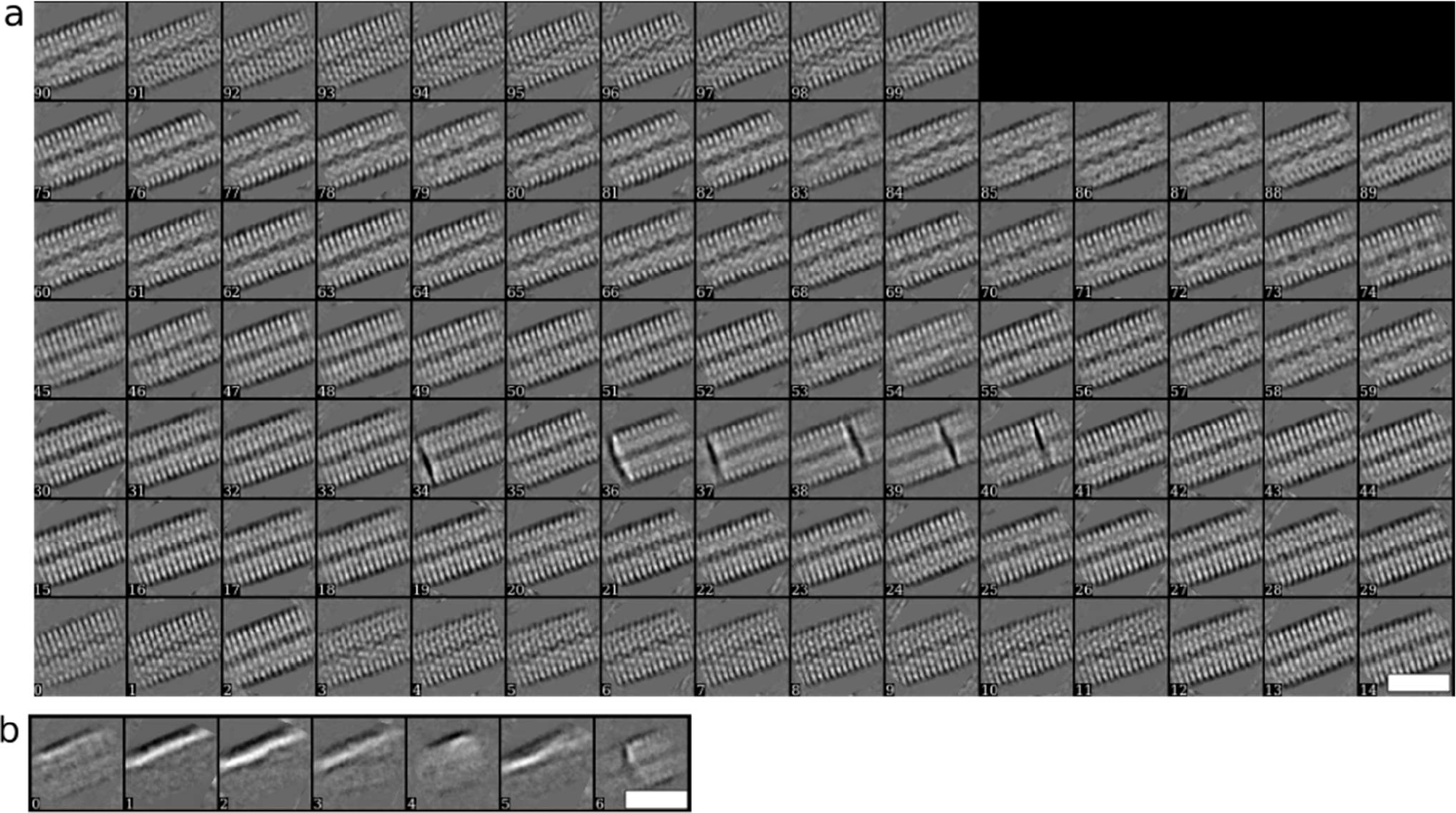
2D classes of TMV accepted or rejected by Cinderella. (a) 100 out of 496 accepted classes and (b) all rejected classes for filaments picked by STRIPER. Scale bars, 25 nm.

**Supplementary Figure 6.**
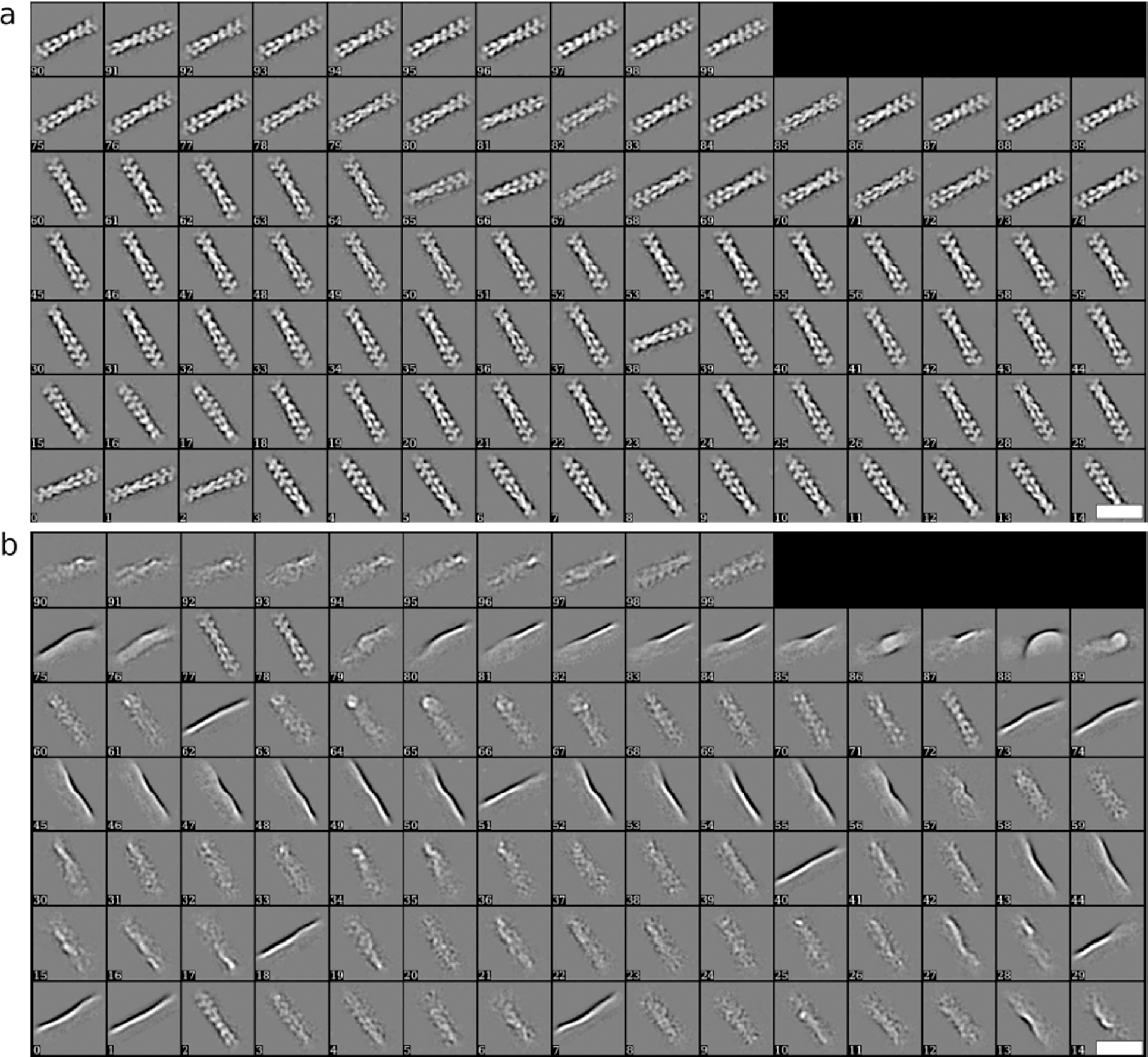
2D classes of F-actin filaments accepted or rejected by Cinderella. (a) 100 out of 708 accepted classes and (b) 100 out of 420 rejected classes for filaments picked by STRIPER. Scale bars, 25 nm.

